# Cell–cell adhesion in plant grafting is facilitated by β-1,4-glucanases

**DOI:** 10.1101/2020.03.26.010744

**Authors:** Michitaka Notaguchi, Ken-ichi Kurotani, Yoshikatsu Sato, Ryo Tabata, Yaichi Kawakatsu, Koji Okayasu, Yu Sawai, Ryo Okada, Masashi Asahina, Yasunori Ichihashi, Ken Shirasu, Takamasa Suzuki, Masaki Niwa, Tetsuya Higashiyama

## Abstract

Plant grafting is conducted for vegetative propagation in plants, whereby a piece of living tissue is attached to another tissue through establishment of cell–cell adhesion. Plant grafting has a long history in agriculture and has been applied to improve crop traits for thousands of years^1^. Plant grafting has mostly relied on the natural ability of a plant for wound healing. However, the compatibility of cell–cell adhesion typically limits graft combinations to closely related species^2–4^, and the mechanism by which cell–cell adhesion of injured tissues is established is largely unknown. Here, we show that a subclade of β-1,4-glucanases secreted into the extracellular region facilitates cell–cell adhesion near the graft interface. *Nicotiana* shows a propensity for cell–cell adhesion with a diverse range of angiosperms, including vegetables, fruit trees, and monocots, in which cell wall reconstruction was promoted in a similar manner to conventional intrafamily grafting^5–7^. Using transcriptomic approaches, we identified a specific clade of β-1,4-glucanases that is upregulated during grafting in successful graft combinations but not in incompatible grafts and precedes graft adhesion in inter- and intrafamily grafts. Grafting was facilitated with an overexpressor of the β-1,4-glucanase and, using *Nicotiana* stem as an interscion, we produced tomato fruits on rootstocks from other plant families. Our results demonstrate that the mechanism of cell–cell adhesion is partly conserved in plants and is a potential target to enhance plant grafting techniques.

---

Plant grafting is the procedure of connecting two or more pieces of living plant tissues to grow as a single plant, for which the healing of the wound site is accomplished through the adhesion of proliferated cells^2–4^. Use of grafting is necessary for propagation of many fruit trees worldwide, such as apples, pears, grapes, and citrus, for vegetable cultivation in Asian and European countries to obtain the benefits of certain rootstocks, such as disease resistance and tolerance of unfavorable soil conditions, and to control the quantity and quality of fruit^1,8^. Recently, grafting has been used in scientific studies to explore the mechanisms of systemic signaling in plants where long-distance transport of phytohormones, RNAs, and proteins in the vascular system has an important molecular basis^9–11^. Although grafting is a useful technique, for practical use the scion–stock combination (the shoot and root parts of a graft, respectively) is limited to closely related plant species. In general, grafting is successful between members of the same species, genus, and family, but not between members of different families because of graft incompatibility^2–4,9^. However, several interfamily graft combinations have been reported^12–17^, including a combination we have studied previously involving a *Nicotiana benthamiana* scion (*Nb*, Solanaceae) and an *Arabidopsis thaliana* stock (*At*, Brassicaceae)^18^, in which the *Nb* scion grew slowly but distinctly (Supplementary Movie 1, Extended data Fig. 1a, b). Moreover, in nature, plants that parasitize species from a different plant family have evolved a haustorium, a specialized organ that invades host plant tissues and absorbs nutrients following tissue adhesion^19^. Therefore, plants potentially have the ability to achieve cell–cell adhesion between members of different families.

**Figure 1.**
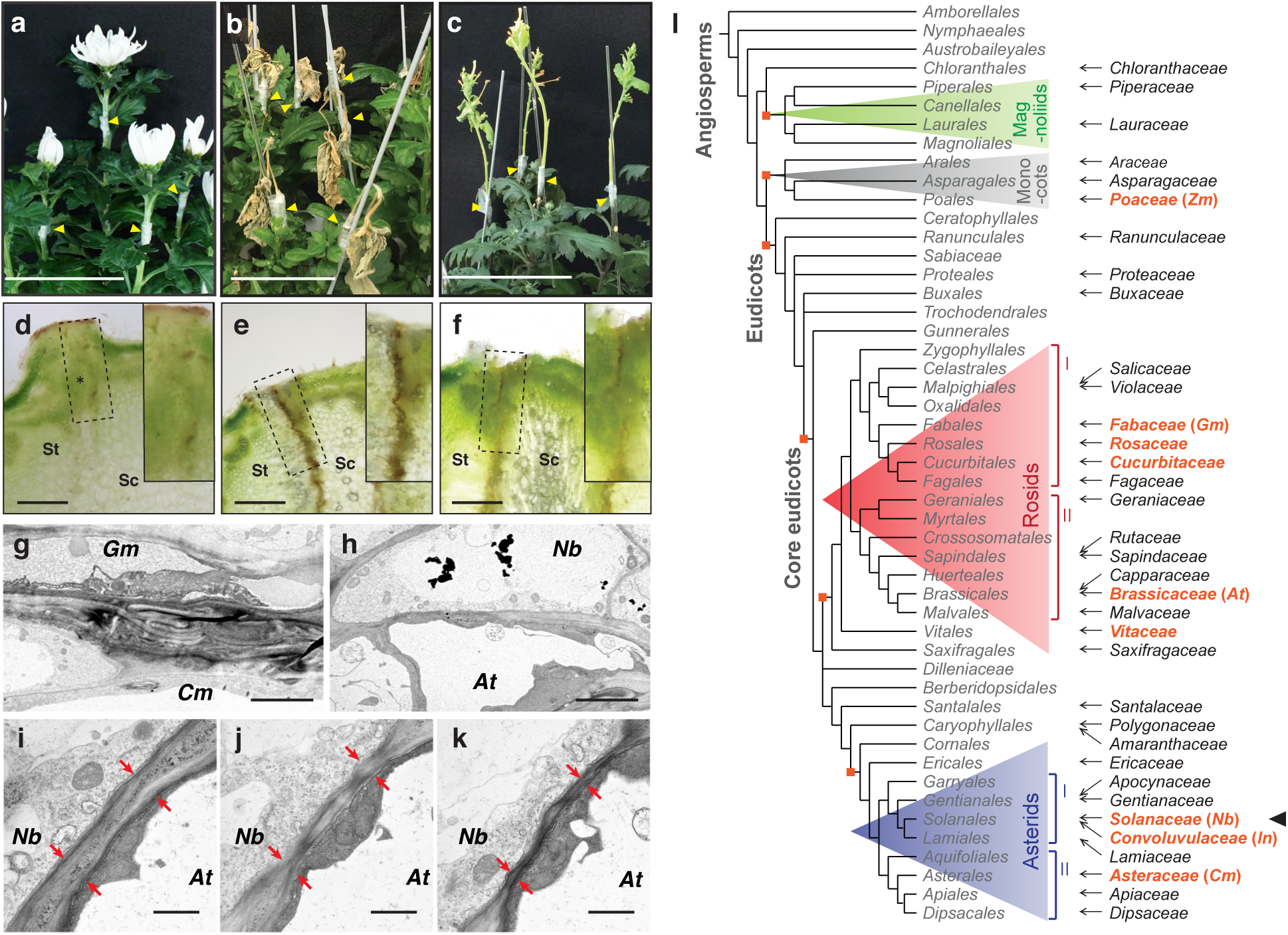
*Nicotiana* established cell–cell adhesion in interfamily grafting. **a**–**c**, Four weeks after grafting of the *Cm, Gm*, and *Nb* scion on the *Cm* stock. Scale bars, 10 cm. **d**–**f**, Transverse sections at graft junctions of (**a**–**c**). Dashed rectangles indicate the position of insets. In the *Gm*/*Cm* interfamily graft, a necrotic layer is formed at the graft interface (**e**), but is less developed in the *Cm*/*Cm* homograft (**d**) and *Nb*/*Cm* interfamily graft (**f**). Scale bars, 1 mm. **g**, Transmission electron micrograph (TEM) near the *Gm*/*Cm* graft junction showing folding of the cell walls. Scale bar, 5 µm. **h**, Stacked cell walls is not observed near the junction of the *Nb*/*At* interfamily graft. Scale bar, 5 µm. **i**–**k**, TEMs for serial sections of a cell–cell boundary at the graft interface of a *Nb*/*At* interfamily graft 2 weeks after grafting. Arrows indicate the thickness of the cell wall between the cells. Scale bars, 1 µm. **l**, Phylogenetic tree showing angiosperm families with which *Nicotiana* species (an arrowhead) form compatible interfamily grafts (arrows). Families including major crops are indicated in red.

## *Nicotiana* potential for interfamily grafts

We expanded the range of graft combinations to include other angiosperms and observed that *Nicotiana* shows strong potential for cell–cell adhesion with phylogenetically distant plant species and interfamily grafts are alive for more than 1 month (Fig. 1, Supplementary Tables 1, 2). In the case where *Chrysanthemum morifolium* (*Cm*, Asteraceae) was used as the stock and we conducted *Cm*/*Cm* homografts (scion/stock notation) and interfamily grafts with *Glycine max* (*Gm*, soybean, Fabaceae) and *Nb*, the *Cm*/*Cm* homografts established and the *Cm* scions produced flowers, whereas the *Gm*/*Cm* interfamily grafts did not establish and the *Gm* scion died (Fig. 1a, b). By contrast, in *Nb*/*Cm* interfamily grafts, the *Nb* scion continued to grow four weeks after grafting (Fig. 1c). The *Nb* scion grew for more than three months until setting seeds. The viability of *Nicotiana* interfamily grafting was confirmed in combinations using *Nb* as the stock (Extended data Fig. 1c). In transverse sections of the graft junctions for these combinations, a necrotic layer formed at the graft boundary in unsuccessful *Gm*/*Cm* interfamily grafts but developed only weakly in successful *Cm*/*Cm* homografts and *Nb*/*Cm* interfamily grafts two weeks after grafting (Fig. 1d–f). Necrotic layer formation is an indicator of incompatibility in cell–cell adhesion in grafting^15–17^. These results indicated that *Nb*/*Cm* grafts showed cell–cell adhesion despite the interfamily combination. Transmission electron microscopy (TEM) revealed folded cell wall remnants caused by graft injury at the graft interface of *Gm*/*Cm* unsuccessful interfamily grafts (Fig. 1g). In contrast, a thin cell wall formed in some areas of the graft interface in *Nb*/*At* interfamily grafts (Fig. 1h). Serial sections indicated a decrease in cell wall thickness at the graft interface (Fig. 1j–k, Extended data Fig. 2). These results are consistent with previous observations of cellular morphology at graft interfaces of compatible interfamily grafts^16^. Taken together, these findings indicate that certain plant species, such as *Nb* and other species studied, can accomplish cell–cell adhesion in interfamily combinations. We then examined how widespread this capability may be among angiosperms. We conducted grafting experiments using plants of seven *Nicotiana* species (*Nb* was predominantly used) and an interfamily partner from 84 species in 42 families, chosen from among 416 angiosperm families^20^. Ability for cell–cell adhesion was evaluated based on scion viability 4 weeks after grafting, because in incompatible combinations scion viability is lost soon after grafting and the loss of viability is visible after transferring grafts to a low-humidity environment. We observed that *Nicotiana* species showed compatibility in interfamily grafting with 73 species from 38 families, including two species of magnoliids, five species of monocots, and 65 species of eudicots, such as important vegetable, flower, and fruit tree crops, with *Nicotiana* plants used as either the scion or stock (Fig. 1l, Extended data Fig. 3, Supplementary Tables 1, 2). These observations indicated that the cell–cell adhesion capability of *Nicotiana* plants could be extended for grafting with a diverse range of angiosperms.

**Figure 2.**
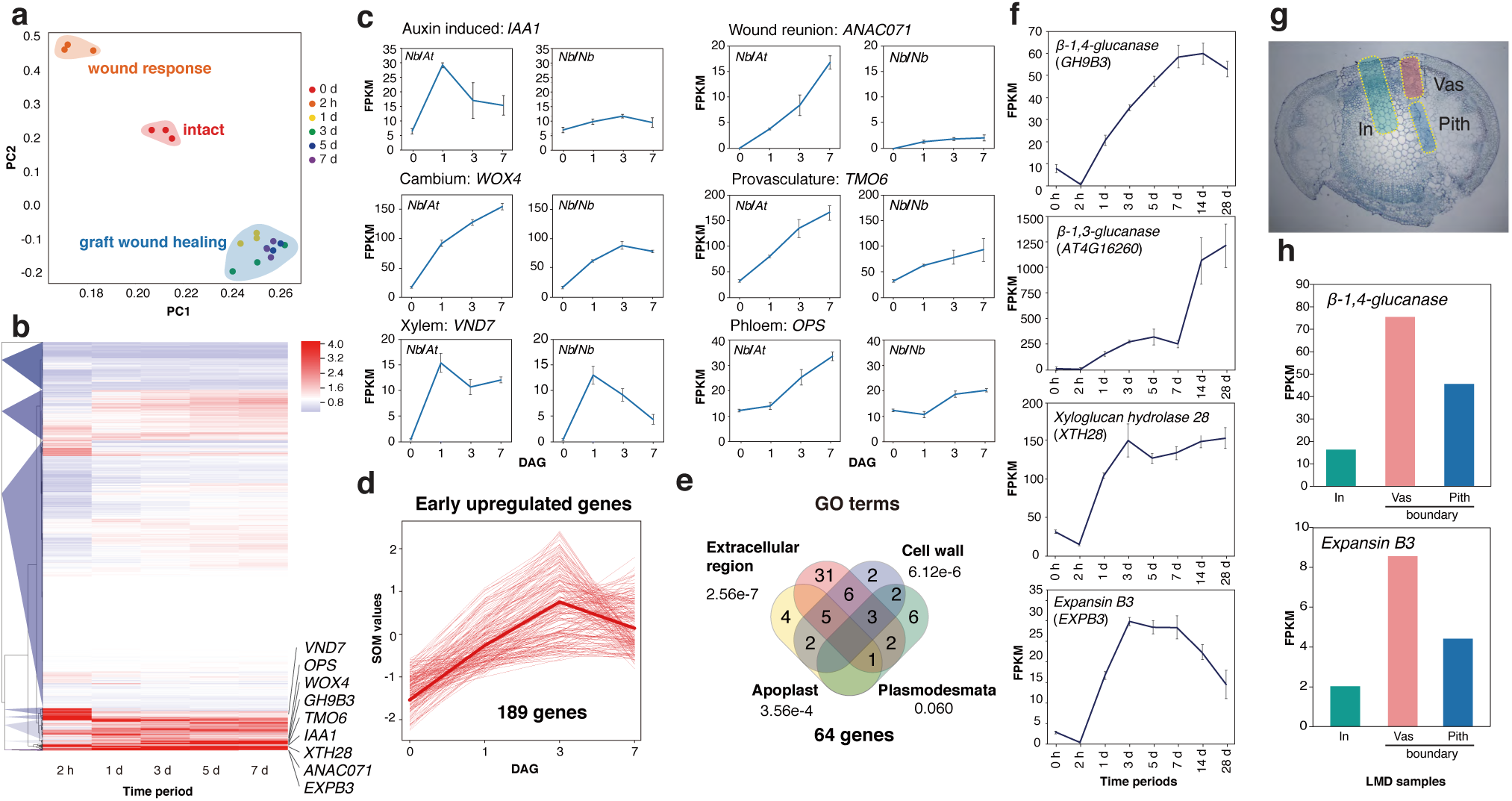
Transcriptomic analysis revealed conventional graft-associated gene expression in *Nicotiana* interfamily grafting. **a**, Principal component analysis of the transcriptome of the *Nb* intact stem and the scion of *Nb*/*At* interfamily grafts at five time points (three biological replicates for each time points) distinguishes intact stems, wound response a short time after grafting, and the graft wound healing process. PC, principal component. **b**, Hierarchical clustering with Euclidean distance and Ward’s minimum variance method over ratio of RNA-seq data from five time points after *Nb*/*At* grafting against intact plants resolves nine gene clusters. Genes associated with grafting reported in previous studies are marked. **c**, Expression levels of genes associated with auxin action, wound repair, and cambium, provascular xylem, and phloem development in *Nb*/*At* interfamily grafts and *Nb*/*Nb* homografts. Supplementary Information Table 3 provides details. **d**, Extraction of early-upregulated genes associated with heterograft formation. Bold line indicates average of 189 genes. **e**, GO enrichment analysis of 189 genes shows enrichment of ‘Extracellular region’, ‘Cell wall’, ‘Apoplast’, and ‘Plasmodesmata’. Genes in the four categories overlap. Each numerical value represents the *P*-value of the GO analysis. **f**, Expression profile of representative genes among the 189 early-upregulated genes after grafting. **g**, Laser microdissection of *Nb*/*At* heterograft tissue was performed for the RNA-seq analysis. In, Vas, and Pith represent the inner central area of *Nb* scion tissue, and the cambial and pith area of the graft boundary for RNA extraction samples, respectively. **h**, LMD-RNA-seq of genes presented in (**f**) shows significant expression in Vas.

**Figure 3.**
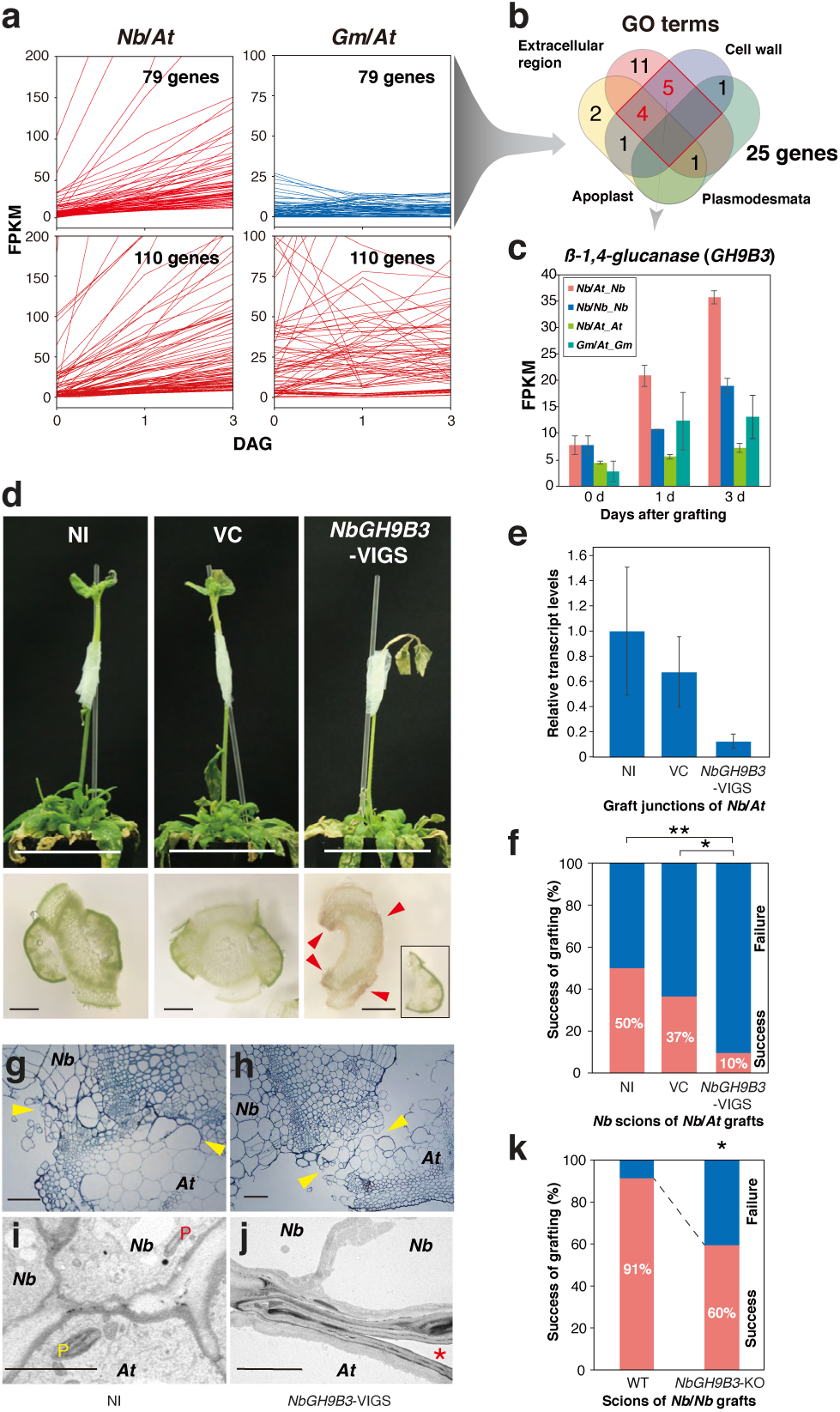
Cell wall modification involved in *Nicotiana* interfamily grafting. **a**, Search for genes involved in *Nb* interfamily grafting. Of 189 upregulated genes in *Nb* (Fig. 2d), 110 genes were upregulated but 79 genes were not in incompatible *Gm*/*At* interfamily grafts. Expression patterns of the 110 and 79 genes in *Nb*/*At* and *Gm*/*At* are shown. **b**, GO enrichment analysis of 79 genes shows the genes overlapping in two categories, ‘Extracellular region’ and ‘Cell wall’, were mostly extracted after classification in (**a**) and are marked in red. **c**, Expression profile of *β-1,4-glucanase* (*NbGH9B3*) in represented samples. **d**, *Nb*/*At* grafts two weeks after grafting in VIGS experiments (upper panels). *Nb* scions infected with CMV virus harboring a partial sequence of *NbGH9B3* to trigger gene silencing (*NbGH9B3*-VIGS), with no virus infection (NI) and vector control (VC) were grafted. Lower panels show transverse sections of each graft junction. Inset indicates the intercept of *At* tissues separated. **e**, Suppression of *NbGH9B3* expression by VIGS was verified by qRT-PCR. Expression levels were normalized against *NbACT1* and adjusted to be relative to the NI sample. **f**, Effect of the *NbGH9B3*-VIGS on graft establishment. Differences between the sample groups were tested with Fisher’s exact tests with α set at *P* < 0.05 (*) or *P* < 0.01 (**), *n* = 30 grafts for each sample fraction. **g**–**j**, Transverse sections of grafted stem sample as represented. **g, h**, Optical microscopic images. Arrowheads indicate the boundary of *Nb* and *At*. Scale bars, 100 µm. **i, j**, TEM images. Yellow and red ‘P’ indicate the plastid of *At* and *Nb*, respectively. * indicates a gap formed between *Nb* and *At* cells. Scale bars, 5 µm. **k**, Effect of CRISPR knock-out (KO) of *NbGH9B3* on graft establishment. The effect of KO was evaluated using Fisher’s exact test (*P* < 0.05). Graft establishment was confirmed for 41 of 45 wild-type grafts and 28 of 47 KO grafts.

## *Nicotiana* promotes cell wall reconstruction during interfamily grafting

To examine the underlying mechanism of *Nicotiana* interfamily grafting, we performed transcriptome analysis on graft junction samples from *Nb*/*At* interfamily grafts 2 h after grafting and 1, 3, 5, 7, 10, 14, and 28 days after grafting (DAG) with the following controls: intact *Nb* stem, and graft junctions of *Nb*/*Nb* homografts at 1, 3, 5, and 7 DAG (Fig. 2). The transcriptome was distinctly changed 2 h after grafting compared with that of intact *Nb* and the transcriptome changed gradually over time after grafting (Fig. 2a, b). On the basis of clustering data, genes previously reported to be associated with grafting^21,22^ (Supplementary Table 3) were upregulated in response to *Nb*/*At* interfamily grafting, including genes associated with auxin action, wound repair, and cambium, provascular and vascular development (Fig. 2b). The expression level was comparable or relatively higher than that observed in *Nb*/*Nb* homografts (Fig. 2c), which may indicate that *Nicotiana* interfamily grafting requires greater contribution of these genes to achieve graft establishment of more weakly compatible combinations. These molecular responses at the graft junction were consistent with morphological changes in the *Nicotiana* interfamily grafts, in which cell proliferation and xylem bridge formation were observed in the grafted region but the xylem bundle was obviously thin (Extended data Fig. 4a–e). Dye tracer experiments using toluidine blue, an apoplastic tracer, and carboxyfluorescein, a symplasmic tracer, provided evidence for establishment of both apoplastic and symplasmic transport at 3 DAG or later (Extended data Fig. 4f–h). Moreover, transport of mRNAs^18^ and GFP proteins across the graft junction was also detected (Extended data Fig. 4i, j), although the amount detected was weaker than that for homografts. Hence, the viability of the *Nb* scions was preserved by parenchymatous tissue formation at the graft interface.

**Figure 4.**
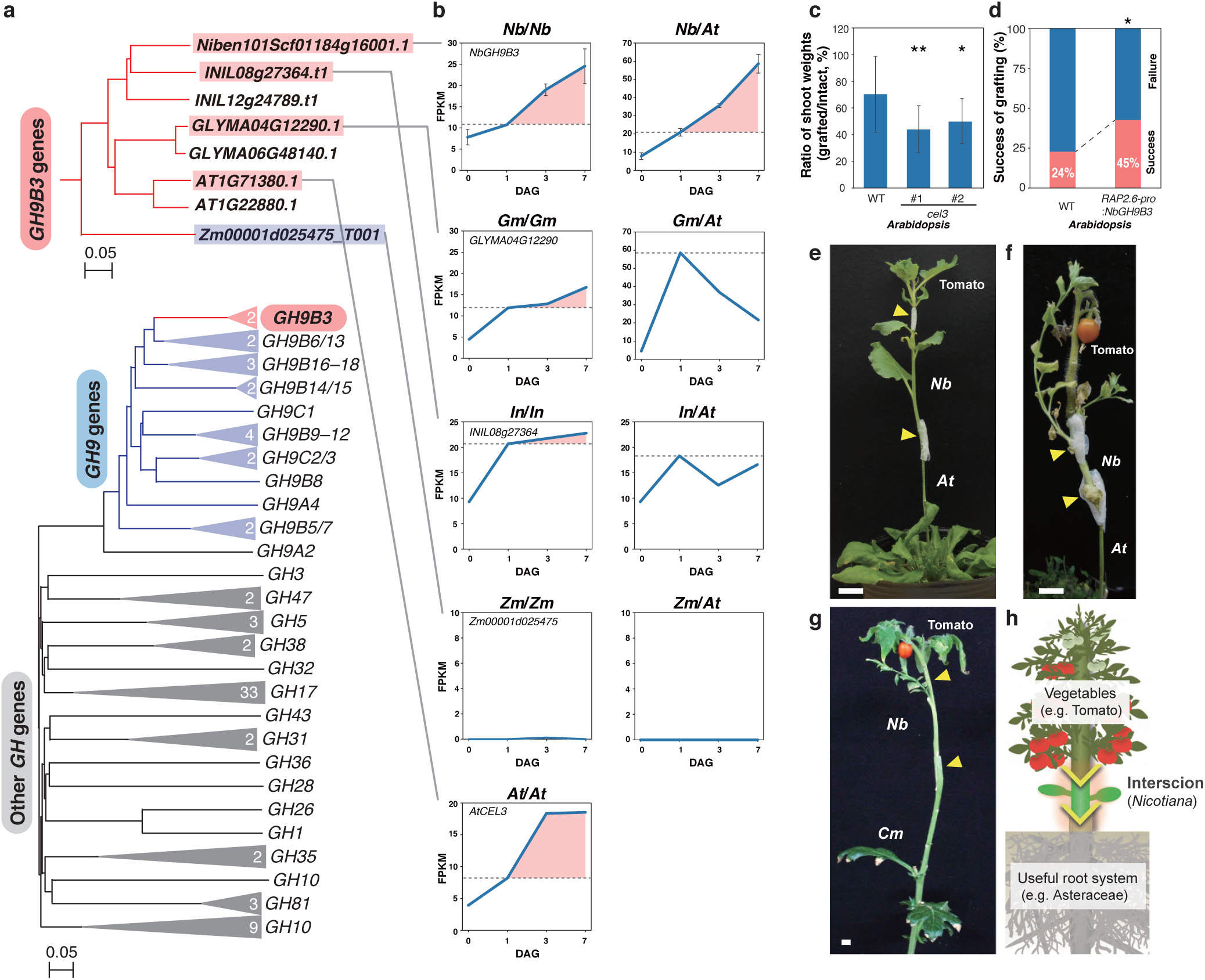
*Glycosyl hydrolase 9B3* is essential for graft wound healing in plants. **a**, Phylogeny of plant glycosyl hydrolase gene family including the *GH9B3* clade. Upper panel shows a tree for the *GH9B3* clade genes and lower panel shows a tree for all *GH* clades. The number of *At* genes included in each clade is shown in triangles (see Methods). **b**, *GH9B3* clade genes located in the same clade as *Niben101Scf01184g16001* show a common expression pattern; expression of genes up-regulated at an early stage after grafting is maintained when the graft is established, or continues to rise subsequently, and is not maintained if the graft is not established. **c**, Increase in shoot fresh weight after grafting in two lines of mutants for *AtCEL3*, a *GH9B3* clade gene in *At*, and the wild type. Experiments were performed on 14–19 seedlings for each sample fraction. Student’s *t-*tests were conducted (\**P* < 0.05). **d**, An *At* overexpression line of *NbGH9B3* using a RAP2.6 wound-inducible promoter (*NbGH9B3*-OX) increased percentage success of grafting compared with wild-type grafting. In the experiment, graft trials were performed on 64 *NbGH9B3*-OX and 102 wild-type seedlings. Viability of the scion was determined two weeks after grafting and the effect of overexpression was evaluated by Fishers’ exact test (*P* < 0.05). **e**–**g** Grafts of tomato scion onto *At* (**e**; 3 weeks after grafting, **f**; 4 months after grafting) or *Cm* (**g**; 3 months after grafting) using a *Nb* interscion. Arrowheads indicate grafted points. Scale bars, 1 cm. **h**, Proposed method to perform interfamily grafting mediated by a *Nicotiana* interscion.

To elucidate the molecular events in the early stage of cell–cell adhesion, we extracted early-upregulated genes in the *Nb* scion of *Nb*/*At* interfamily grafts and identified 189 genes (Fig. 2d, Supplementary Table 3). In a gene ontology (GO) enrichment analysis for these genes, the top-ranked GO terms were ‘Extracellular region’, ‘Cell wall’ and ‘Apoplast’ (Fig. 2e, Methods), which indicated that cell wall modification was undertaken in *Nicotiana* interfamily grafting. Genes encoding cell wall modification/reconstruction enzymes, including β-1,4-glucanase, β-1,3-glucanase, xyloglucan hydrolase, and expansin, were promoted at 1 to 28 DAG (Fig. 2f). Laser microdissection samples of *Nb*/*At* interfamily graft junctions confirmed the enhanced expression level of a number of these genes in the cells proliferated from the cambial or pith region of the graft boundary (Fig. 2g, h). Expression of genes associated with cell wall dynamics was also implicated in previous transcriptomic studies of conventional intrafamily grafting^5–7^ and wounding response^23^, which implies that *Nicotiana* activates a mechanism for cell wall reconstruction in either intra- or interfamily grafting.

## *Nicotiana* interfamily grafting requires a secreted type of β-1,4-glucanase

We next investigated the characteristics of *Nicotiana* grafting by comparing the transcriptome with that of interfamily grafting using soybean (*Gm*), which was incompatible in interfamily grafting combinations. We screened genes that were upregulated in the *Nb* scions of *Nb*/*At* interfamily grafts but not in the *Gm* scions of *Gm*/*At* interfamily grafts. For comparisons, we selected each homologous gene in *Gm* that showed the highest homology using tblastx (see Methods). Using the gene information obtained, we extracted genes that were upregulated in *Nb* scions but not in *Gm* scions. Of 189 genes upregulated in *Nb* scions (Fig. 2d), upregulation of 79 genes was not observed in *Gm* scions (Fig. 3a, Methods). We assumed that these genes may explain the difference in graft compatibility of *Nb* and *Gm*. Among the 79 genes, genes associated with ‘Extracellular region’ and ‘Cell wall’ were highly conserved (Fig. 3b) (in comparison with Fig. 2e, the number of genes associated with ‘Extracellular region’ and ‘Cell wall’ was nine out of 14, whereas the number of genes associated with the other GO terms was 16 out of 50). This result again suggested that cell wall reconstruction is a critical event for success of interfamily grafting of *Nicotiana*. This population included a gene encoding β-1,4-glucanase of the glycosyl hydrolase 9B (GH9B) family, designated *NbGH9B3* based on similarity to *At* genes. Expression of *NbGH9B3* was significantly upregulated in the *Nb*/*At* interfamily grafts in contrast to corresponding genes in the *At* stocks of *Nb*/*At* and the *Gm* scions of *Gm*/*At* interfamily grafts (Fig. 3c). Given that β-1,4-glucanases of the GH9B family show cellulolytic activities and play roles in cellulose digestion, relaxation of cell wall, and cell wall construction during plant growth processes, such as root elongation^24,25^, we hypothesized that *NbGH9B3* facilitates cell– cell adhesion of opposing cells at the graft boundary and further analyzed *NbGH9B3* function in grafting.

We applied virus-induced gene silencing (VIGS) to examine the function of *NbGH9B3* in *Nb*/*At* interfamily grafting (Fig. 3d–f). We prepared non-infected and vector control samples for comparison. VIGS targeting of *NbGH9B3* caused failure of *Nb*/*At* interfamily grafting 2 weeks after grafting; the *Nb* scion was easily detached from the *At* stocks and the *Nb* tissues formed a necrotic layer on the graft surface (Fig. 3d). The expression level of *NbGH9B3* was consistent with the percentage success of grafting (Fig. 3e, f). At the graft interface of *Nb* scions in which *NbGH9B3* was down-regulated by VIGS, folded cell walls were frequently observed in contrast to the non-infected control (Fig. 3g–j). We generated a knock-out line of *NbGH9B3* (*NbGH9B3*-KO) using a clustered, regularly interspaced, short palindromic repeats (CRISPR)/CRISPR-associated protein 9 nuclease (CRISPR/Cas9) editing method (see Methods) and conducted grafting experiments. The percentage success of grafting wild-type *Nb* scions onto *At* stocks was 91%, whereas that of *NbGH9B3*-KO interfamily grafting was 60% (Fig. 3k). These data suggested that the β-1,4-glucanase encoded by *NbGH9B3* functions in cell wall digestion at the graft interface and facilitates graft establishment in *Nicotiana* interfamily grafting.

## GH9B3 plays a crucial role in graft establishment

We hypothesized that *Nicotiana* interfamily grafting with a diverse range of plants is achieved through a common mechanism of cell–cell adhesion during graft formation. To test this hypothesis, we examined whether β-1,4-glucanase also functions in conventional intrafamily grafting for other genera (Fig. 4a, b). We prepared homograft samples of soybean (*Gm*), morning glory (*In*), maize (*Zm*), and *Arabidopsis* (*At*). Among *GH9* family genes, one gene from each plant species was distinctly upregulated at 1 to 7 DAG in all homografts except maize homografts, which failed to graft successfully because monocot species lack cambial activity in the stem^26^; the genes all belonged to the *GH9B3* clade (Fig. 4a, b, Extended data Fig. 5). Moreover, the *GH9B3* genes were temporally upregulated at 1 DAG but expression was not increased subsequently in *Gm* and *In* scions grafted onto *At* stocks (Fig. 4b). These data suggested that upregulation of *GH9B3* genes during graft adhesion was conserved among these plants and, in the case of *Nicotiana*, this mechanism can be switched on even in interfamily grafting. In *Zm* grafts, an orthologous gene was not upregulated in both homografts and interfamily grafts, which may imply that during the evolution of maize the mechanism to promote expression of *GH9B3* clade genes in response to stem injury was lost.

To examine the role of *GH9B3* genes in grafting in other plant genera, we performed seedling micrografting in *Arabidopsis* using wild-type and two T-DNA insertion mutant lines for *AtCEL3*, a *GH9B3* clade gene that was upregulated in *At* homografts (Fig. 4b). Although a significant difference in percentage success was not observed among wild-type and mutant homografts, shoot growth after grafting was significantly decreased in grafts of both mutant lines compared with that of the wild type (Fig. 4c), which indicated that GH9B3 is required for straightforward establishment of the graft connection in *At* and that dependency on GH9B3 is higher in *Nicotiana* interfamily grafting (Fig. 3f, k). We next examined the effect of GH9B3 overexpression on grafting. We generated transgenic lines of *Arabidopsis* that overexpressed *NbGH9B3* under the control of a wound-induced *RAP2*.*6* promoter (*NbGH9B3*-OX)^27^. The percentage success of micrografting using the *NbGH9B3-*OX line was significantly higher than that of wild-type grafting (Fig. 4d). Thus, it was demonstrated that GH9B3 functions in graft formation in plants other than *Nicotiana*.

The aforementioned results indicated that *Nicotiana* plants activate a mechanism for graft adhesion even in interfamily grafting, which is generally activated only in the case of intrafamily healable grafting. The ubiquity of GH9B3, an enzyme secreted into the extracellular region, in plants enables the success of *Nicotiana* interfamily grafting with a diverse range of angiosperms. To exploit this capability, we examined whether *Nicotiana* could act as an intermediate in the grafting of different plant families. We chose tomato as the scion because the fruit exhibit several favorable traits, such as umami flavor, nutrient richness, and lycopene production, and are cultivated worldwide^28^. We grafted the tomato scion onto *At* or *Cm* stocks using a *Nicotiana* interscion, where the junction between the tomato scion and the *Nicotiana* interscion represented an intrafamily graft. The tomato scions were successfully stabilized and ultimately produced fruit 3–4 months after grafting (Fig. 4e–g, Extended data Fig. 6a). We also achieved other interfamily grafts in which the scion, interscion, and stock all belonged to different plant families (Supplementary Table 4, Extended data Fig. 6b). One of the stock plants we used was *Cm*, a member of the Asteraceae, one of largest family in angiosperm, which is economically important family for oils and leaf vegetables such as sunflower seeds and lettuce, and at the same time, is recognized as invasive weeds in various circumstances including non-arable region^29^. Our results therefore demonstrated that grafting might increase the utilization of beneficial root systems found in natural resources with minimal destruction of ecosystems (Fig. 4h).

Grafting is reliant on plants’ ability for tissue adhesion and healing of wounds, which is fundamental for the hardiness and vigor of plants in nature. Grafting is achieved through sequential cellular processes, including wound response, cell regeneration, cell proliferation, cell–cell adhesion, and cell differentiation into specific tissues, and is an important topic in plant science^9–12^. In regard to cell–cell adhesion, the cell wall polysaccharide matrix is heterogeneous and varies considerably in composition among plant species, therefore in interspecific grafting differences in cell wall composition may account for incompatibility. *Nicotiana* shows cell–cell adhesion compatibility with diverse plant species through the function of a conserved clade of extracellular-localized β-1,4-glucanases, the GH9B3 family, which probably target cellulose, a core structural component of the cell wall in plants^24,30^, together with other components associated with cell wall dynamics (Fig. 2). Thus, we identified a typical biological function of a specific clade of glycosyl hydrolase large gene family (hundreds number of genes are included in the family, Fig 4a) through a study on grafting. Cellular processes involved in grafting require characterization and the outcomes of such studies may enhance grafting techniques in plant science research and agriculture worldwide.

## Methods

### Plant materials

*Nicotiana benthamiana* seeds were surface sterilized with 5% (w/v) bleach for 5 min, washed three times with sterile water, incubated at 4°C for 3 days, and sown on half-strength Murashige and Skoog (1/2 MS) medium supplemented with 0.5% (m/v) sucrose and 1% agar. The pH was adjusted to pH 5.8 with 1 M KOH. Seedlings were grown at 22°C for *At* and 27°C for *Nb* under continuous illumination of 100 µmol m^−2^ s^−1^. For generation of the *Nb NbGH9B3*-KO lines, three artificially synthesized DNA fragments (5′-TGTCAAGTTCAATTTCCCAA-3′, 5′-CAATGCTTTCTTGGAACACA-3′, and 5′-CCGATTACTTCCTCAAGTGT-3′) were cloned as sgRNA into the pTTK352 vector for CRISPR/Cas9 editing^31^. Binary vectors were introduced into *Agrobacterium tumefaciens* strain EHA105 by electroporation and transformed into *Nb* plants by the leaf disk transformation method^32^. For generation of the *At NbGH9B3*-OX lines, a 1687 bp promoter sequence of *RAP2*.*6* (At1g43160) and a 3078 bp cDNA sequence of *Niben101Scf01184g16001* amplified separately by PCR from the *RAP2*.*6* plasmid vector and *Nb* cDNA library, respectively, were cloned into pENTR/D-TOPO (Thermo Fisher Scientific, Waltham, MA, USA) using InFusion^®^ (Takara Bio, Ohtsu, Japan) as an entry clone. The entry clone was transferred into the pGWB1 vector^33^ using the LR reaction. The sequences of the primers used for PCR amplification were as follows: gw_RAP2.6pro_F2, 5′-CACCTCTAGATGGGATGGTGTACTACGGATG-3′ and gw_RAP2.6pro_R2, 5′-GATCGGGGAAATTCGGTACCCCTCTAGGTTTGAAATTGCGGTGGTAG-3′ for the RAP2.6 promoter; RAP2.6_01184g16001_F, 5′-TTCAAACCTAGAGGGATGGCGTTTAGAGTGAAAG-3′and pgwb_01184g16001_R, 5′-GATCGGGGAAATTCGCTAACGTTTGGAACTATCAA-3′ for the *Niben101Scf01184g16001* CDS. Binary vectors were introduced into *Agrobacterium tumefaciens* strain GV3101 by electroporation. Plant transformation was performed using the floral dip method^34^. The *At cel3* T-DNA insertion lines SALK_032323 and CS_803355 were obtained from the Arabidopsis Biological Resource Center (ABRC; Ohio State University, Columbus, OH, USA). The *At* ecotype Columbia (Col) was used as the wild type.

### Grafting

For the grafting of *Nb* as the scion, 1-to 2-month-old *Nb* inflorescence stems were used. For the other graft combinations, plants of sufficient size to perform grafting were used. For the majority of herbaceous species, 2-week-old to several-month-old plants were used and, for tree species, several-year-old plants were used. Wedge grafting was performed on the epicotyls, stems, petioles, or peduncles. For stock preparation, stems (or other organs) were cut with a 2–3 cm slit at the top. For scion preparation, the stem was cut and trimmed into a V-shape. The scion was inserted into the slit of the stock and wrapped tightly with parafilm. A plastic bar was set along the stock and the scion for support. The entire scion was covered with a plastic bag, which had been sprayed inside with water beforehand. Grafted plants were grown for 7 days in an incubator at 27°C under continuous light (∼30 µmol m^−2^ s^−1^), or in a greenhouse at 22–35°C under natural light (500–1500 µmol m^−2^ s^−1^ during the day). After this period, the plastic bags were partly opened by cutting the bags and making holes for acclimation. The next day, the plastic bags were removed and the grafted plants were grown in an incubator at 22–25°C under continuous light (∼100 µmol m^−2^ s^−1^), in a plant growth room at 22–30°C under continuous light (∼80 µmol m^−2^ s^−1^), or in a greenhouse. Grafting was determined to be successful if the scion was alive at 4 weeks post-grafting for the combinations listed in Supplemental Tables 1–3. For the test of *GH9B* loss- and gain-of-function effect on a percentage of grafting success (Fig. 3f, k, Fig. 4d) was evaluated based on scion viability 2 weeks after grafting. When three plants were grafted using an interscion, grafting manipulations were performed either all at once or in two steps. In the latter case, two graft combinations were performed first; in one case, the future stock was grafted with a *Nb* scion and, in the other case, the future scion was grafted onto a *Nb* stock. After establishment of each graft, a second grafting was performed using the *Nb* parts of each graft. To compare watering and grafting (Extended data Fig. 1b), *Nb* primary stems of 7 cm length were cut and the cut edge was trimmed into a V-shape. Expanded leaves (more than 1 cm width) were removed so that water absorption was directed to stem growth dependent on the cutting sites. Half of the trimmed *Nb* stems were watered only and the other half were grafted as scions onto *At* stocks. The stem length was measured once per week after these treatments (*n* = 24 per treatment). All other plant materials for stem grafting used in this study are listed in Supplemental Tables 1–3.

Micrografting of *At* was performed using a microscaled device constructed for micrografting following a protocol described previously^35^. Seeds were sown on the devices and the devices were put on the Hybond-N+ nylon membrane (GE Healthcare, Chicago, IL, USA) which was placed on 1/2 MS medium supplemented with 0.5% sucrose and 1% agar. Four-day-old seedlings underwent micrografting on the device. After grafting, the device containing grafted seedlings were transferred to fresh 1/2 MS medium supplemented with 0.5% sucrose and 2% agar and grown at 22°C (for the test of *NbGH9B3*-OX line) or 27°C (for the test of *At cel3* mutant lines) for 6 days. After this period, the grafted seedlings were taken out from the device and transferred to fresh 1/2 MS medium supplemented with 0.5% sucrose and 1% agar and grown at 22°C. The phenotype was examined at 10 DAG.

### Microscopy

To capture brightfield images of hand-cut sections of grafted regions, a stereomicroscope (SZ61, Olympus, Tokyo, Japan) equipped with a digital camera (DP21, Olympus) or an on-axis zoom microscope (Axio Zoom.V16, Zeiss, Göttingen, Germany) equipped with a digital camera (AxioCam MRc, Zeiss) was used.

To observe xylem tissues, hand-cut transverse sections of the grafted stem region were stained with 0.5% toluidine blue or 1% phloroglucinol. Phloroglucinol staining (Wiesner reaction) was performed using 18 µL of 1% phloroglucinol in 70% ethanol followed by addition of 100 µL of 5 N hydrogen chloride to the section samples. Brightfield images were captured using a stereomicroscope or a fluorescence imaging microscope.

To determine apoplasmic transport, the stems of *At* stocks were cut, and the cut edge was soaked in 0.5% toluidine blue solution for 4 h to overnight. Hand-cut transverse sections of the grafted regions were observed using a stereomicroscope (SZ61, Olympus) or a fluorescence imaging microscope (BX53, Olympus) equipped with a digital camera (DP73, Olympus) for high-magnification images. The water absorption sites were stained blue.

To determine symplasmic transport, cut leaves from *At* stocks were treated with 0.01% 5(6)-carboxyfluorescein diacetate (CF; stock solution 50 mg ml^−1^ in acetone), together with 0.1% propidium iodide (PI) to distinguish symplasmic transport (indicated by CF fluorescence) from apoplastic transport (indicated by PI fluorescence) for 4 h to overnight. Transverse sections of the grafted regions and the apical regions of *Nb* scions were hand-cut and observed. To examine GFP protein transport, *Nb* scions were grafted onto transgenic *35S::EGFP At* stocks ^36^. Hand-cut transverse sections of the grafted regions were made and the fluorescence images were captured using a fluorescence imaging microscope (BX53, Olympus) or a confocal laser scanning microscopy (LSM780-DUO-NLO, Zeiss). To quantify the GFP fluorescence signal in the grafted region, lambda mode scanning was performed by collecting emissions in the 490–658 nm range with excitation at 488 nm and extracted against the GFP reference spectrum. The tile scan mode was also used to capture a wide view of the entire graft section. The *z*-sectioning images were processed using ZEN 2010 software to create maximum-intensity projection images.

To observe resin-embedded sections, samples were fixed with 2% paraformaldehyde and 2% glutaraldehyde in 0.05 M cacodylate buffer (pH 7.4) at 4°C overnight. After fixation, the samples were washed three times with 0.05 M cacodylate buffer for 30 min each, and then postfixed with 2% osmium tetroxide in 0.05 M cacodylate buffer at 4°C for 3 h. The samples were dehydrated in a graded ethanol series (50%, 70%, 90%, and 100%). The dehydration schedule was as follows: 50% and 70% for 30 min each at 4°C, 90% for 30 min at room temperature, and four changes of 100% for 30 min each at room temperature. Dehydration of the samples was continued in 100% ethanol at room temperature overnight. The samples were infiltrated with propylene oxide (PO) two times for 30 min each and transferred to a 70:30 mixture of PO and resin (Quetol-651, Nisshin EM Co., Tokyo, Japan) for 1 h. Then, the caps of the tubes were opened and PO was volatilized overnight. The samples were transferred to fresh 100% resin and polymerized at 60°C for 48 h. For light microscopy, the polymerized samples were sectioned (8 µm thickness) with a microtome and mounted on glass slides. For light microscopic observation, sections of 1.5 µm thickness were stained with 0.5% toluidine blue (pH 7.0), mounted on the glass slides with Mount-Quick (Daido Sangyo Co., Tokyo, Japan), and observed using a digital microscope (DMBA310, Shimadzu RIKA Co., Tokyo, Japan). For transmission electron microscopic analysis, the polymerized samples were ultra-thin-sectioned at 80–120 nm with a diamond knife using an ultramicrotome (ULTRACUT UCT, Leica, Tokyo, Japan). The sections were mounted on copper grids and stained with 2% uranyl acetate at room temperature for 15 min, then washed with distilled water followed by secondary staining with lead stain solution (Sigma-Aldrich Co., Tokyo, Japan) at room temperature for 3 min. The grids were observed using a transmission electron microscope (JEM-1400Plus, JEOL Ltd, Tokyo, Japan) at an acceleration voltage of 80 kV. Digital images were captured with a CCD camera (VELETA, Olympus Soft Imaging Solutions GmbH, Münster, Germany).

### Transcriptome analysis

The grafted or intact plants were harvested at the respective time points. Approximately 10–15 mm of graft junction or stem tissue at a similar location was sampled. Each biological replicate comprised the pooled tissues from 10 grafts or 10 intact plants. Total RNA was extracted from the samples using the RNeasy Mini Kit (Qiagen, Hilden, Germany) following the manufacturer’s protocol. The cDNA libraries were prepared with an Illumina TruSeq Stranded Total RNA kit with Ribo-Zero Plant or the BrAD-Seq method^37,38^ and sequenced for 86 bp single end with an Illumina NextSeq 500 platform (Illumina, San Diego, CA, USA). Data preprocessing was performed as follows. Raw sequence quality was assessed with FastQC v0.11.4 (http://www.bioinformatics.babraham.ac.uk/projects/fastqc/). Adapters were removed and data trimmed for quality using Trimmomatic v0.36 with the settings TruSeq3-PE-2.fa: 2:40:15, SLIDINGWINDOW: 4:15, LEADING: 20, TRAILING: 20, and MINLEN: 30 (http://www.usadellab.org/cms/). FastQC quality control was repeated to ensure no technical artifacts were introduced. Trimmed reads were mapped on the genome assembly using HISAT2 version 2.1.0 with the settings -q -x “$index” –dta —dta-cufflink (http://daehwankimlab.github.io/hisat2/). The generated SAM files were converted to BAM format and merged using SAMtools version 1.4.1 (http://samtools.sourceforge.net). Gene expression levels (fragments per kilobase of transcript per million fragments mapped; FPKM) were estimated using Cufflinks version 2.1.1 with the -G option (http://cole-trapnell-lab.github.io/cufflinks/). The expression fluctuation profiles were generated using Cuffdiff version 2.1.1. The reference sequences and version used for mapping and annotation were as follows: *Nb*,https://btiscience.org/our-research/research-facilities/research-resources/nicotiana-benthamiana/, *Nicotiana benthamiana* draft genome sequence v1.0.1; *At*, https://www.arabidopsis.org, TAIR10 genome release; *Gm*, https://phytozome.jgi.doe.gov/pz/, Phytozome v7.0 (Gmax_109); *In*, http://viewer.shigen.info/asagao/,Asagao_1.2; and *Zm*, https://plants.ensembl.org/Zea_mays/, Zea_mays.AGPv4.

Extraction of upregulated genes (Fig. 2d) was performed using the Cuffdiff results of *Nb*/*At* and *Gm*/*At* graft samples, with three biological replicates for each, according to the following criteria: (i) for evaluation using the ratio between the two samples, genes whose expression level (FPKM value) is >0 in the 0 DAG samples, (ii) the ratio of the value at 1 DAG to that at 0 DAG is 2 or more, (iii) the value at 3 DAG is higher than that at 1 DAG, (iv) the values at 5 and 7 DAG are higher than that at 0 DAG, and (v) the value at 1 DAG is higher than 10. Based on the *Nb* transcript sequence of the extracted 189 genes, homology analysis of the amino acid sequence was performed with tblastx using the *At* transcript as a reference, and the *At* gene ID closest to each gene was obtained. A GO enrichment analysis was performed with DAVID (https://david.ncifcrf.gov) using the obtained *At* gene IDs and a Venn diagram was created. For each of the 189 genes, homology analysis on *Gm* was performed using tblastx to obtain the orthologous gene of *Gm*. Data classification for Fig. 3a was performed according to the following criteria: (i) the FPKM values at 1 and 3 DAG were lower than 15, and (ii) the ratio at 3 DAG to 1 DAG was 1.5 or less. For construction of phylogenetic tree for plant glycosyl hydrolase gene family (Fig. 4a), *GH9B3* clade genes were isolated from *Nb, Gm, In, Zm*, and *At* as well as the other *GH* genes from *At* on the Phytozome database (http://www.phytozome.net) or the TAIR database (https://www.arabidopsis.org) and were used to test the phylogeny. For the *At* genes, we handled 88 genes which harbor *GH* numbers on the annotations. The tree shown in Fig. 4a was reconstructed with a part of the entire phylogenetic tree using the neighbor-joining method. Upper panel shows a tree topology for the *GH9B3* clade genes of *Nb, Gm, In, Zm*, and *At*. Lower panel shows a tree for all *At GH* genes where only the primary branches for each *GH* groups were drawn. The number of genes included in each clade is shown in triangles. The *GH9B3* clade we called includes *AtGH9B3* and *AtGH9B4*. Data extraction for Fig. 4b was performed using the Cuffdiff results of *Nb*/*At* samples as described above and *Gm*/*Gm, Gm*/*At, In*/*In, In*/*At, Zm*/*Zm, Zm*/*At*, and *At*/*At* graft samples in biological replicates for each tissue at each time point.

For laser microdissection (LMD) samples, stem segments of the graft junction of *Nb*/*At* interfamily grafts (∼15 mm) were frozen in liquid nitrogen. Frozen samples were embedded with Super Cryoembedding Medium (Section-Lab, Hiroshima, Japan) in a dry ice/hexane cooling bath, and then cryosectioned into 15-µm-thick transverse sections using a cryostat (CM1860, Leica) in accordance with the method of Kawamoto ^39^. The sections that adhered to films were desiccated in a −20°C cryostat chamber for 30–60 min. Sections of three tissue regions from heterografts (vascular tissue adjacent to the graft union, pith tissue adjacent to the graft union, and *Nb* scion tissue) were microdissected using a LMD6500 laser microdissection system (Leica) and separately collected into RNA extraction buffer composed of buffer RLT (Qiagen) and 0.01% β-mercaptoethanol. Total RNA was extracted using a QIAshredder (Qiagen) and the RNeasy Plant Mini kit (Qiagen) in accordance with the manufacturer’s instructions. The cDNA libraries were constructed using an Ovation RNA-Seq System V2 (NuGEN Technologies, Redwood City, CA, USA) in accordance with the manufacturer’s instructions. RNA sequencing (RNA-Seq) analysis was performed as described above. RNA-Seq data are available from the DNA Data Bank of Japan (DDBJ; http://www.ddbj.nig.ac.jp/).

### VIGS experiments

For VIGS of *Niben101Scf01184g16001* (*NbGH9B3*), a 294 bp portion of the region spanning from the 5′ UTR to the first exon of *NbGH9B*3 was amplified by PCR using primers harboring a partial sequence of *NbGH9B*3 and CMV-A1 vector: GA_F_NbGH9B3, 5′-GTCACCCGAGCCTGAGGCCTGAAAAAGACACTTGATCGAAAAGC-3′; and GA_R_NbGH9B3, 5′-GGGGAGGTTTACGTACACGCGTGTCCTTCAAAGAACAAAATGG-3′. For the no-silencing vector control, a 292 bp portion of the *GFP* gene was amplified by PCR usingthefollowingprimers:GA_F_GFP, 5′-GTCACCCGAGCCTGAGGCCTGACTCGTGACCACCCTGACCTAC-3′;and GA_R_GFP, 5′-GGGGAGGTTTACGTACACGCGGCTTGTCGGCCATGATATAGA-3′.The amplified fragments were cloned between the *Stu*I and *Mlu*I sites of the CMV-A1 vector using the Gibson assembly method^40^. Plasmids containing the full-length cDNA of the viral RNA were transcribed *in vitro* and leaves of 3-week-old *Nb* plants were dusted with carborundum and rub-inoculated with the transcripts. Successful infection of the virus in the upper leaves of *Nb* plants without deletion of the inserted sequences was confirmed by RT-PCR of the viral RNA using the primers CMV_RNA2_2327_F, 5′-ATTCAGATCGTCGTCAGTGC-3′, and CMV_RNA2_2814_R, 5′-AGCAATACTGCCAACTCAGC-3′. Primary inoculated leaves were used for secondary inoculation of *Nb* plants. Primary inoculated leaves were ground in 100 mM phosphate buffer (pH 7.0) and 10 µL of the homogenate was placed dropwise on the three expanded leaves of a new 3-week-old *Nb* plant and rub-inoculated. One week after inoculation (corresponding to age 4 weeks), the infected *Nb* stem was grafted onto the bolting stem of 5-week-old *At* plants. Two weeks after grafting, the percentage success of grafting was scored based on scion survival.

Quantitative reverse-transcription PCR was performed using cDNA templates prepared from total RNA from the stem of intact *Nb* or grafted plants 3 days after grafting. The PCR conditions were 50°C for 2 min, 95°C for 10 min, and 40 cycles of 95°C for 15 s followed by 60°C for 1 min; The primers used were qPCR_Cellulase5_F2, 5′-ATTGGGAGCCAATGATGTACC-3′, and qPCR_Cellulase5_R2, 5′-TGTCATTTCCAACAACGCTTC-3′. *NbACT1* (*Niben101Scf09133g02006*.*1*) was used as the internal standard and amplified with the primers NbACT-F, 5′-GGCCAATCGAGAAAAGATGAC-3′, and NbACT-R, 5′-AACTGTGTGGCTGACACCATC-3′. All experiments were performed with three independent biological replicates and three technical replicates.

## Acknowledgements

We thank D. Kurihara, T. Araki, K. Shiratake, S. Otagaki, S. Ishiguro, and Japan Tobacco Inc., Japan for plant materials and A. Iwase for the *RAP2*.*6* plasmid vector. We thank T. Shinagawa, H. Fukada, A Ishiwata, R. Masuda, Y. Hakamada, M. Hattori, M. Matsumoto, I. Yoshikawa, A. Yagi, A. Shibata, and A. Furuta for technical assistance, and T. Akagi and Y. Hattori for discussions. This work was supported by grants from the Japan Society for the Promotion of Science Grants-in-Aid for Scientific Research (18KT0040, 18H03950 and 19H05361 to M.N. and 2516H06280 and 19H05364 to Y.S.), the Cannon Foundation (R17-0070), the Project of the NARO Bio-oriented Technology Research Advancement Institution (Research Program on Development of Innovative Technology 28001A and 28001AB) to M.N., and the Japan Science and Technology Agency (ERATO JPMJER1004 to T.H. and START15657559 and PRESTO15665754 to M.N.).

## Author Contributions

M.N., K.K., Y.S., and M. Niwa conceived of the research and designed experiments. M.N., and K.O. performed grafting experiments. M.N., and Y. Sawai analyzed tissue sections. M.N. collected microscopic data with Y.S.’s support. R.T. performed VIGS experiments. Y.K. performed micrografting experiments. M.A. collected LMD samples. K.K., R.O., and Y.I. generated RNA-Seq libraries. T.S. performed sequencing. K.K. analyzed transcriptome data. M.N., M. Niwa, K.S., and T.H. supervised the experiments. M.N., K.K. and M. Niwa wrote the paper.

