## Supplemental file for "Cell–cell adhesion in plant grafting is facilitated by β-1,4-glucanases"

**Extended Data Figure 1. Growth of *Nicotiana* interfamily grafts.**

**Extended Data Figure 2. Cell walls in close contact at the graft boundary of *Nb*/*At* interfamily graft.**

**Extended Data Figure 3. *Nicotiana* interfamily grafts with a diverse range of angiosperms.**

**Extended Data Figure 4. *Nicotiana* interfamily grafts established apoplastic and symplasmic transport.**

**Extended Data Figure 5. Expression patterns of GH9B family genes.**

**Extended Data Figure 6. *Nicotiana* interscion mediated successful interfamily grafting.**

**Supplementary Tables 1. Grafting experiments in which *Nicotiana* was used as the scion.**

**Supplementary Tables 2. Grafting experiments in which *Nicotiana* was used as the stock.**

**Supplementary Tables 3. Gene annotation list for Fig. 2.**

**Supplementary Tables 4. Upregulated genes in the *Nb*/*At* interfamily grafts.**

**Supplementary Tables 5. Grafting experiments in which *Nicotiana* was used as the interscion.**

**Supplementary Video 1. Growth of a *Nb*/*At* interfamily graft at 8 to 32 DAG.**

*Author for correspondence: Michitaka Notaguchi

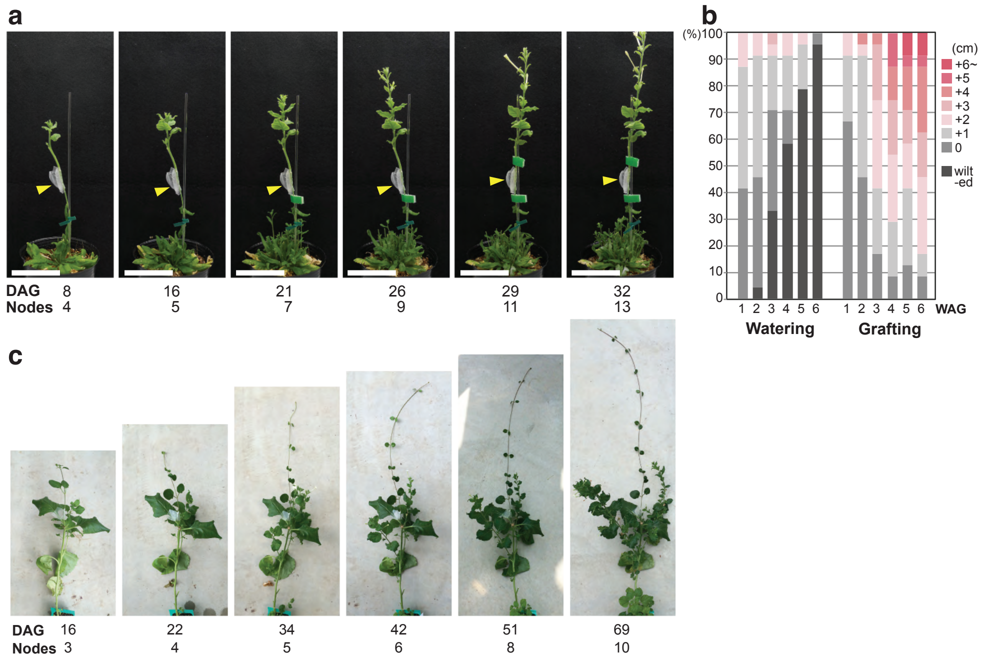


**Extended Data Figure 1. Growth of *Nicotiana* interfamily grafts.**
**a**, Time series of the growth of a *Nicotiana benthamiana* (*Nb*) scion grafted onto an *At* stock. Days after grafting (DAG) and the number of nodes produced on the scions (Nodes) are indicated below the images. Arrowheads indicate graft points. Scale bars, 5 cm. **b**, Growth of *Nb* cut stems soaked in water (Watering) or grafted onto *At* stock (Grafting). Stem growth (cm) measured every week after grafting (WAG) is indicated. **c**, Growth of *Nicotiana* interfamily grafts where *Nb*, as the stock, was grafted with a *Vinca major* (*Vm*) scion. Age of grafted plants (DAG) and the number of nodes produced on the scions (Nodes) are indicated below each image. Arrowheads indicate grafted points. Scale bars, 5 cm.

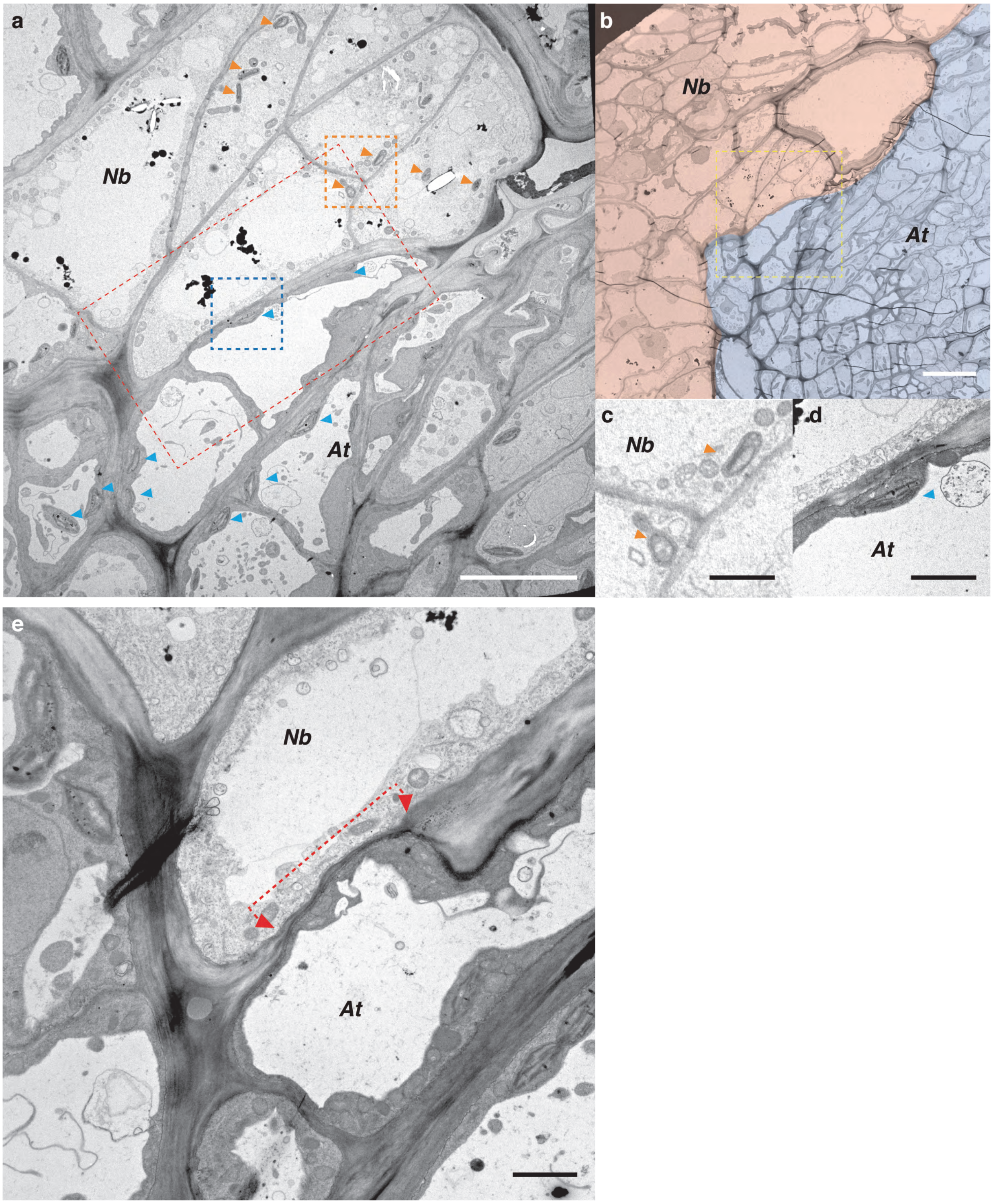


**Extended Data Figure 2. Cell walls in close contact at the graft boundary of *Nb*/*At* interfamily graft.**
**a**, Transmission electron micrograph near the graft boundary between *Nb* and *At*. Red dashed rectangle corresponds to Fig. 1h. *Nb* and *At* cells can be distinguished from the shape of the plastid. Orange and blue arrowheads indicate *Nb* and *At* plastids, respectively. Orange and blue dashed rectangles indicate the area of **c** and **d**, respectively. Scale bar, 10 µm. **b**, Based on the shape of plastids, the boundary between *Nb* and *At* is represented by color coding. The pink area is *Nb* and the blue area is *At*. The yellow dashed rectangle corresponds to **a**. Scale bar, 20 µm. **c**, High-magnification image of *Nb* plastids. Scale bar, 2 µm. **d**, High-magnification image of an *At* plastid. Scale bar, 2 µm. **e**, Section corresponding to Fig. 1k. In the area indicated by the dashed arrow, the cell walls of *Nb* and *At* adhere closely without gaps. Scale bar, 2 µm.

a

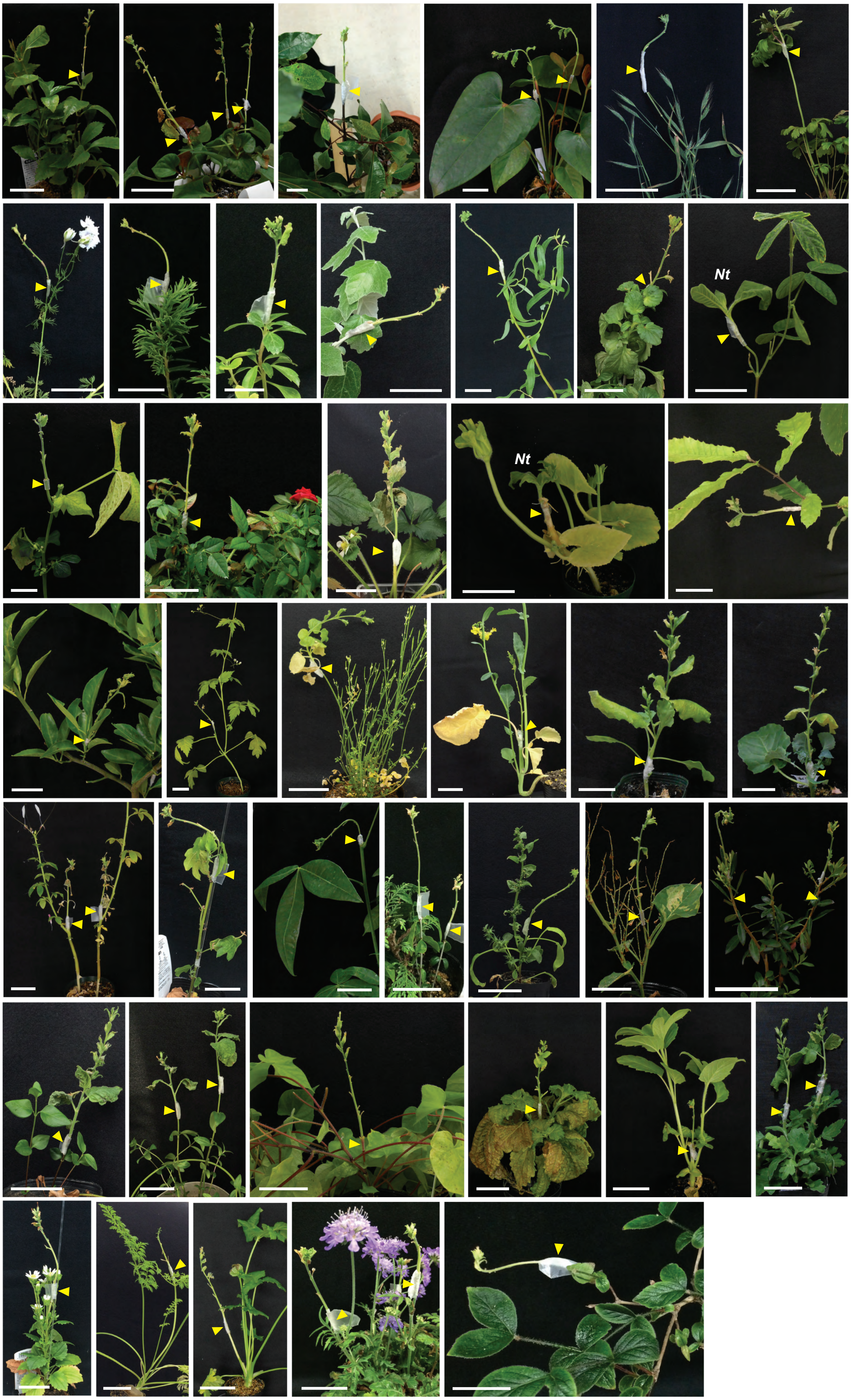


**Extended Data Figure 3.** ***Nicotiana*** **interfamily grafts with a diverse range of angiosperms (Continued).**

**b**

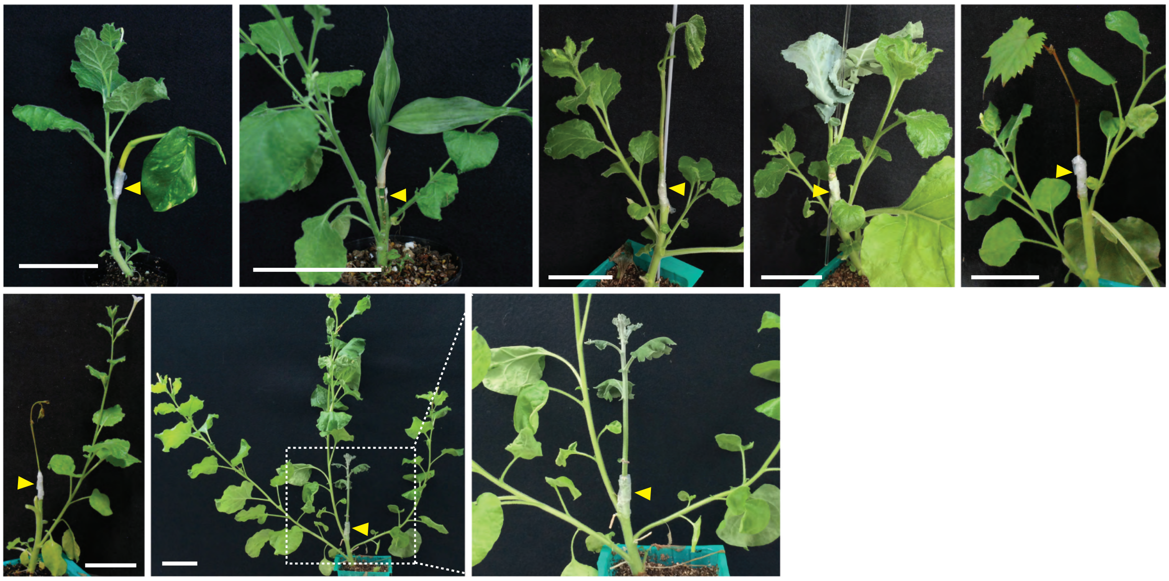

**Extended Data Figure 3. *Nicotiana* interfamily grafts with a diverse range of angiosperms.**
**a**, *Nicotiana* interfamily grafting where *Nb* or *N. tabacum* (*Nt*, marked in the photos) was grafted as the scion onto a series of plants. In the top row (from left to right; plant species used as the stock and the date of the image taken are indicated), *Sarcandra glabra* at 23 DAG, *Houttuynia cordata* at 23 DAG, *Cinnamomum camphora* at 33 DAG, *Anthurium* at 22 DAG, *Brachypodium distachyon* at 28 DAG, and *Anemone coronaria* at 29 DAG; second row, *Consolida* at 37 DAG, *Grevillea* at 33 DAG, *Pachysandra terminalis* at 33 DAG, *Populus alba* at 42 DAG, *Salix matsudana* at 42 DAG, *Viola* × *wittrockiana* at 30 DAG, and *Glycine max* at 42 DAG; third row, *Vigna angularis* at 26 DAG, *Rosa* at 23 DAG, *Cucurbita maxima* at 61 DAG, *Fragaria* at 30 DAG, and *Quercus crispula* at 33 DAG; fourth row, *Citrus unshiu* at 22 DAG, *Cardiospermum halicacabum* at 59 DAG, *Cardamine hirsuta* at 28 DAG, *Brassica napus* at 24 DAG, *Brassica oleracea* var. *capitata* at 29 DAG, and *Brassica oleracea* var. *italica* at 29 DAG; fifth row, *Cleome hassleriana* at 49 DAG, *Gossypium* at 40 DAG, *Pachira* at 22 DAG, *Buckleya lanceolata* at 33 DAG, *Spinacia oleracea* at 30 DAG, *Fallopia japonica* at 23 DAG, and *Rhododendron* at 23 DAG; sixth row, *Vinca major* at 29 DAG, *Gentiana scabra* at 29 DAG, *Ipomoea nil* at 26 DAG, *Perilla frutescens* at 25 DAG, and *Sesamum indicum* at 39 DAG; seventh row, *Chrysanthemum seticuspe* at 29 DAG, *Callistephus chinensis* at 40 DAG, *Daucus carota* at 43 DAG, *Cryptotaenia japonica* at 33 DAG, *Scabiosa atropurpurea* at 33 DAG, and *Abelia spathulata* at 33 DAG. **b**, *Nicotiana* interfamily grafting where *Nb* was grafted as the stock with scion from diverse plant species. Top row (left to right), *Epipremnum aureum* at 28 DAG, *Dracaena sanderiana* at 49 DAG, *Cucumis sativus* at 63 DAG, *Brassica oleracea* var. *italica* at 56 DAG, and a leaf of *Vitis coignetiae* at 29 DAG; second row, a stem bud of *V. coignetiae* at 29 DAG, and *Chrysanthemum seticuspe* at 62 DAG (a magnified image of a marked area is shown in the left). Arrowheads indicate graft points. Scale bars, 5 cm.

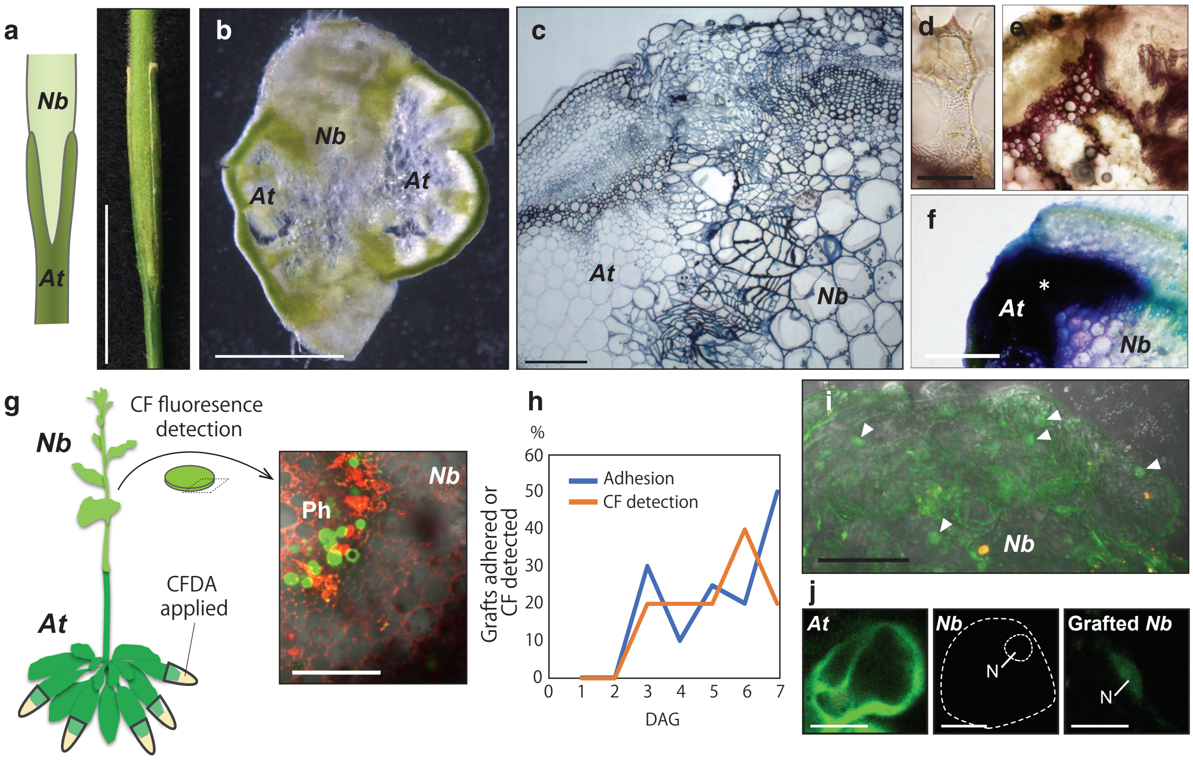


**Extended Data Figure 4. *Nicotiana* interfamily grafts established apoplastic and symplasmic transport.**
**a**–**c**, Morphological characteristics of graft sites of a *Nb* scion grafted onto an *At* stock grown for 2 weeks after grafting. **a**, Wedge-grafted sites (left, a schematic illustration; right, an optical image). **b**, **c**, Hand-cut (**b**) and resin-embedded (**c**) transverse sections of the graft junction region. **d**, **e**, Differentiated xylem tissues in parenchymatous callus tissue proliferated from the *Nb* scions at the graft junction region. **d**, A tracheary element. **e**, Phloroglucinol-stained transverse section showing local xylem bridges between the stock and scion (arrows). **f**, Transverse section of the *Nb*/*At* graft junction after toluidine blue was applied to the *At* stock 2 weeks after grafting. Toluidine blue dye was detected in the *Nb* scion across the graft junction (*), indicating apoplastic transport was established. Scale bar, 500 µm. **g**, **h**, Test of symplasmic transport using carboxyfluorescein diacetate (CF). **g**, Schematic illustration (left) showing that a mixture of CF and propidium iodide (PI), which are symplasmic and apoplastic tracer dyes, respectively, was applied to the leaves of an *At* stock and observed in the *Nb* scion. In a transverse section of the *Nb* scion stem, green fluorescence of CF was detected in the inner phloem tissues, whilst red fluorescence of PI was detected in different patterns. Ph, phloem. Scale bar, 100 µm. **h**, Graft adhesion and symplasmic transport started from 3 days after grafting (10 plants per time point). **i**, GFP fluorescence was detected in the *Nb* callus tissues at the graft junction proliferated from a *Nb* scion grafted onto a *35S::GFP At* stock plant. Nucleus-localized GFP signals are marked (arrowheads). **j**, Spectral analyses of cells in the *At* stock, the intact *Nb* stem and the *Nb* scion in which the GFP signal was detected in the nucleus (N).

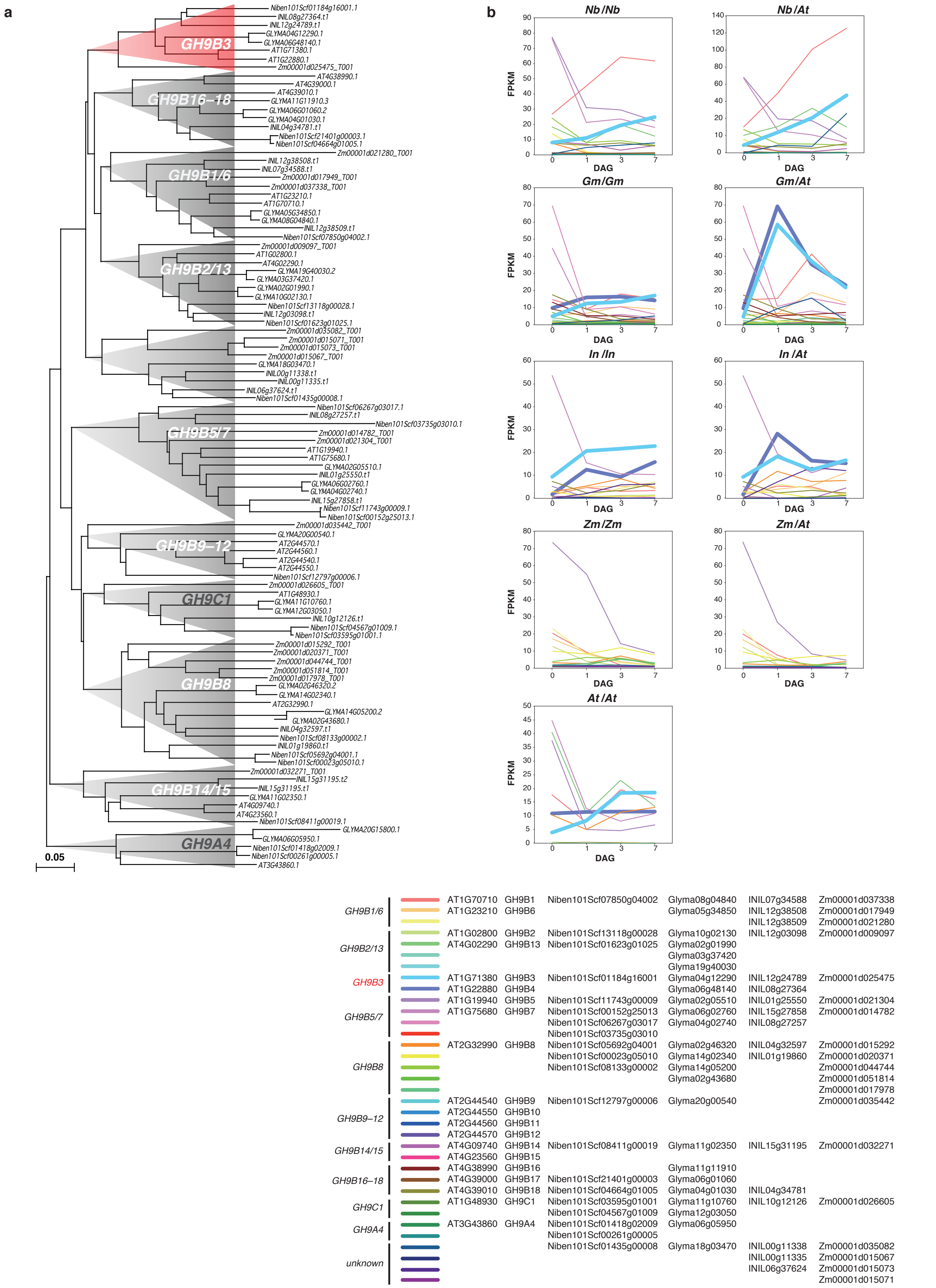


**Extended Data Figure 5. Expression patterns of GH9B family genes.**
**a**, Phylogeny of plant glycosyl hydrolase 9B with some of other glycosyl hydrolase family genes reconstructed from amino acid sequences. **b**, Expression patterns of genes contained in a phylogenetic tree of the GH9B family shown in (**a**). The most similar genes are shown in the same color plots among plant species. The genes contained in the *GH9B3* clade (drawn with thick lines) show similar behavior in the homografts of *Gm* and *In*. Genes belonging to the *GH9B1/6* clade also show increased expression in *Nb* and *Gm*, but not in *In* and *At*. *Zm* shows no increase in expression of any of the genes analyzed.

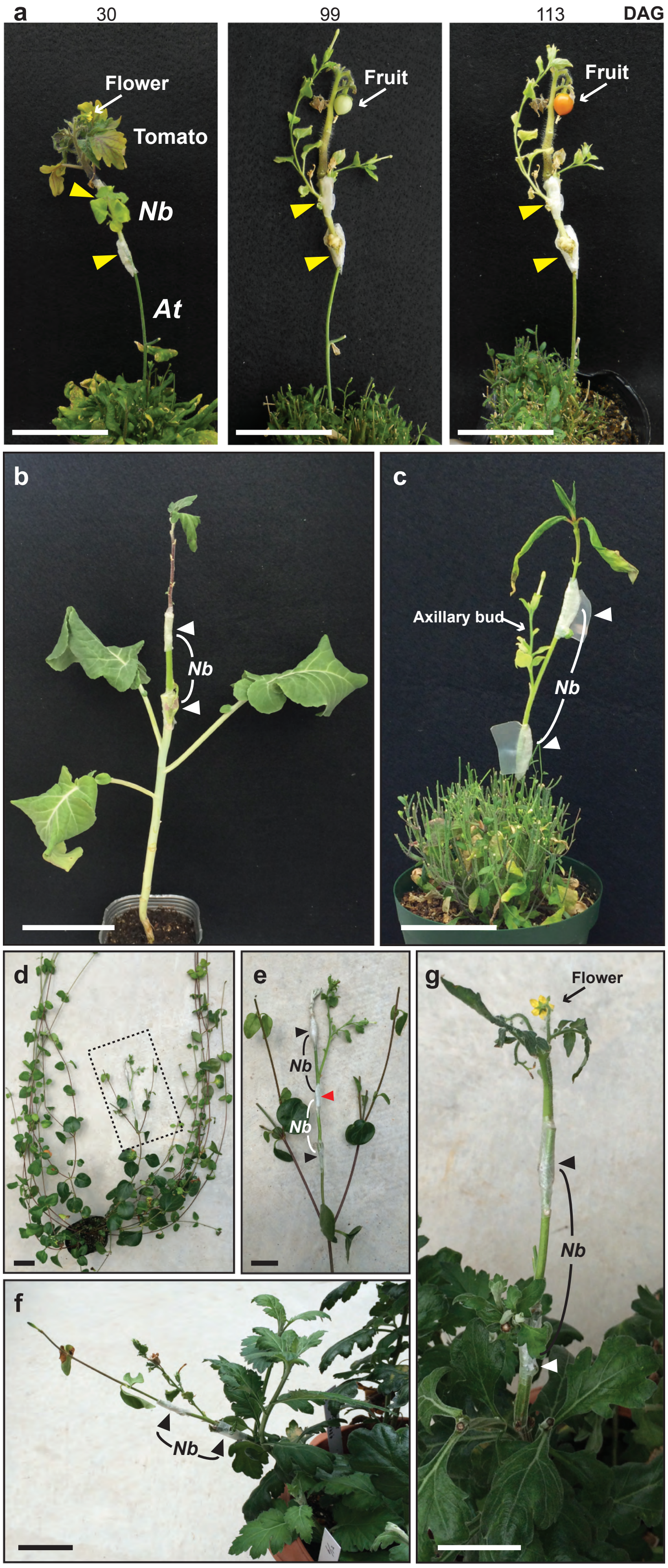


**Extended Data Figure 6. *Nicotiana* interscion mediated successful interfamily grafting.**
**a**, Growth of a *Nicotiana* interfamily graft in which a tomato scion was grafted onto an *At* stock mediated by a *Nb* interscion (identical to Fig. 4f). Age of grafted plants (DAG) is indicated above each photograph. Arrowheads indicate grafted points. Scale bars, 5 cm. **b**–**g**, Photographs of interfamily grafts mediated by *Nb* interscions 1 month after grafting; *Cm* on broccoli (**b**), *Vinca major* on *At* (**c**), *Cm* on *V. major* (**d**, **e**), *V. major* on *Cm* (**f**), and tomato on *Cm* (**g**). Two graft sites were assembled at the same time for the combinations of **c**, **f**, and **g**. For the combination in **b**, the graft of broccoli stock with the *Nb* scion was established first, and the second grafting was performed 28 days after the first grafting, when *Cm* was grafted onto the *Nb* scion. For the combination in **d** (and **e**; a magnified image of the marked area in **d**), two graft combinations were performed first: *Cm* scion/*Nb* stock and *Nb* scion/*V. major* stock (black arrowheads in **e**). The second grafting on the *Nb* scion was performed 28 days after the first grafting (**e**, red arrowhead). Arrowheads indicate the graft points. Scale bars, 5 cm (**b**–**d**, **f**), 1 cm (**e**, **g**).
